# Interplay of primary sequence and RNA secondary structure in determining 5′ splice site choice

**DOI:** 10.1101/376343

**Authors:** Frances Anne Tosto, Asaf Shilo, Jason W. Rausch, Stuart F. J. Le Grice, Tom Misteli

## Abstract

Selective use of 5′ splice sites is a common mechanism by which pre-mRNAs are alternatively spliced. Whereas the sequence requirements of 5′ splice site choice have been well characterized, other important determinants remain poorly defined. Here we apply a combination of structural mapping by SHAPE-MaP and targeted mutational analysis in a cell-based system to comprehensively probe the interplay of primary sequence, secondary RNA structure, regulatory elements and linear splice site position to determine mechanisms of splice site choice *in vivo*. Using the disease-causing alternative 5′ splice site selection in *LMNA* in the premature aging disorder Hutchinson-Gilford Progeria Syndrome as a model system, we identify RNA secondary structural elements near the alternative 5′ splice sites. We show that splice site choice is significantly influenced by the structural context of the available splice sites. While local structure alone is not sufficient to account for splice site selection, the choice of 5′ splice sites depends on the structural stability of the 5′ splice site region which is conferred by downstream elements. In addition, relative positioning of the competing sites within the primary sequence of the pre-mRNA is a predictor of 5′ splice site usage, with the distal position favored over the proximal, regardless of sequence composition. Together, these results reveal an intricate interplay amongst RNA sequence, secondary structure and splice site position in determining 5′ splice site choice.

## Introduction

Our understanding of how RNA impacts diverse cellular processes has grown considerably in recent years. A long appreciated, but still relatively poorly understood regulatory mechanism of RNA function is alternative pre-mRNA splicing^1^, which enables generation of multiple mature mRNAs from the same gene^2^, and as such contributes significantly to proteomic diversity in higher eukaryotes^3^. Critical to alternative splicing is the selection of 5′ and 3′ splice sites (SS).

The strength of a 5′ SS, and thus its likelihood of use, has conventionally been strongly related to its potential for base pairing to the 5′ end of the U1 small nuclear RNA (snRNA)^4^, required for 5′ SS recognition, thus placing much emphasis on the sequence composition of potential 5′ SS. However, 5′ SS selection is now recognized as being a much more complex process involving a number of additional factors that regulate splice site usage. Prominent among these are the proteins of the Ser/Arg-rich (SR)^5–9^ and heterogeneous nuclear ribonucleoprotein (hnRNP) families^10^. In addition, features inherent to the RNA itself have been implicated in 5′ SS choice, including the relative positions of candidate splice sites in the RNA primary sequence^10–13^, as well as local RNA secondary structure, which can suppress 5′ SS usage by limiting accessibility to the spliceosome machinery^14–16^. One notable example of the latter regulatory mechanism is the splicing of *SMN2*, in which a stem-loop structure at the 5′ SS of exon 7 prevents its base pairing with U1 snRNA^17^. While it is clear that multiple factors contribute to splice site choice, how these various determinants are coordinated has not been systematically probed.

Elucidation of the mechanisms of 5′ SS selection has important implications for alternative splicing both in normal physiological processes and in disease since an estimated 10% of disease-causing mutations impact either a 3′ or 5′ SS^18^. One prominent example of disease-related 5′ SS usage is the *de novo* point mutation (C1824T) in exon 11 of the *LMNA* gene that causes the rare premature aging disease Hutchinson-Gilford Progeria Syndrome (HGPS)^19^. The mutation activates an alternative exonic 5′ SS, which in turn results in an internal 150 nucleotide deletion in the mRNA (Δ150) and synthesis of progerin, the disease-causing protein^20^. While slightly higher sequence similarity of the mutant alternative 5′ SS to the consensus 5′ SS sequence likely contributes to its preferential use in HGPS^19^, sequence alone is not sufficient to explain the dramatic increase in alternative 5′ SS usage of mutant LMNA. When assessed by sequence-based, standard splice site scoring algorithms (eg. MaxENT^21^), the alternative 5′ SS score is lower than the nearly perfect score of the normal 5′ SS, even with the point mutation^22^. Based on these considerations, it is clear that non-sequence related factors must contribute to LMNA 5′ SS choice, making the HGPS 5′ alternative splicing event in LMNA an ideal model system to identify determinants of 5′ SS choice.

Here we take advantage of the presence of the two competing 5′ SS in LMNA RNA and perform a structure-function analysis to comprehensively explore the mechanisms of 5′ SS choice. We use SHAPE-MaP to gain detailed structural information on the local and global features of the target RNA and probe the contribution of structural features to 5′ SS choice by systematic mutational analysis using a cell-based system. We find that 5′ SS choice is determined by an intricate interplay of primary sequence, RNA secondary structure, distal regulatory elements and splice site position.

## Materials and Methods

### Cell culture and transfection conditions

HEK293T cells were cultured in Dulbecco’s Modified Eagle Medium (Life Technologies) supplemented with 10% FBS (Sigma), 2 mM L-Glutamine, 1 mM sodium pyruvate, non-essential amino acids, 100 U/ml penicillin and 100 μg/ml streptomycin at 37° C and 5% CO_2_. Cells were seeded in 6 well plates at a density of 350,000 cells/well and incubated overnight. PEI was used for transient transfection of cells with 1μg of plasmid DNA. Following transfection, cells were cultured for an additional 72 hours.

CRL 1474 normal human fibroblast cells were used for antisense oligonucleotide (ASO) experiments. Cells were cultured in Minimum Essential Medium (Life Technologies) supplemented with 15% FBS (Sigma), 2 mM L-Glutamine, 1 mM sodium pyruvate, non-essential amino acids, 100 U/ml penicillin and 100 μg/ml streptomycin at 37° C and 5% CO_2_. Cells were seeded in 60 mm plates at a density of 200,000 cells/plate and incubated overnight. The next day, cells were transfected with the appropriate ASO using Lipofectamine 2000 (Invitrogen) according to the manufacturer’s instructions. Following transfection, cells were cultured for an additional 72 hours. ASO 1919 was purchased from Integrated DNA Technologies and contains 2′-O-Methyl RNA bases with a phosphorothioate backbone. ASO 074 and ASO scrambled (SCR) were obtained from Ionis Pharmaceuticals and contain a 2′-O-methoxy-ethyl ribose modification and a phosphorothioate backbone. ASO sequences are listed in **Supplementary Table 1**.

### Plasmids

LMNA-GFP (WT in this study) and C1824U LMNA-GFP (MUT in this study) minigene splicing reporters were constructed as previously described^23^. A schematic representation of the minigene reporter is shown in **Supplementary Figure 1**. Although not pictured, dsRed is also included in the minigene construct and is used as a loading control. Systematic mutational analysis was performed using the Q5 Site-Directed Mutagenesis Kit (New England BioLabs) based on the manufacturer’s instructions. Primers for site directed mutagenesis (SDM) are listed in **Supplementary Table 2**. The WT/1927A➔G, WT/1927A➔U, WT/Alt-SS-Open-1, WT/1928-1930C➔U, WT/Norm-SS-Closed-1, WT/Norm-SS-Closed-2, and WT/Norm-SS-Sil mutants were all generated from the WT minigene construct. The MUT/1928-1930C➔G, 2X-NORM, MUT/2X-ALT, MUT/Alt-SS-Closed, and MUT/Norm-SS-Open mutants were all generated from the MUT minigene construct. The WT/Alt-SS-Open-2 mutant was generated from the WT/Alt-SS-Open-1 construct. The WT/2X-ALT mutant was generated via two rounds of SDM. In the first round, the MUT/2X-ALT forward and reverse primers were used to make an A➔G substitution at position 1972 of the WT construct. In the second round, the end product of the first round was used as the starting material and the WT/2X-ALT forward and reverse primers were used to introduce a T➔C substitution at position 1974. The SWAP mutant was generated from the MUT/2X-ALT construct. The MUT/Alt-Closed-Norm-Open mutant was generated from the MUT/Alt-SS-Closed construct.

### Secondary structure mapping

Secondary structure models were obtained by SHAPE-MaP structure probing of *in vitro* transcribed segments of WT and C1824U LMNA RNA. Transcription templates containing five 3′-terminal nt of exon 9, all of exons 10 and 11, and all but two nt of intron 11/12 were amplified by PCR from the LMNA-GFP and C1824U LMNA-GFP expression vectors described above. The two 687 nt RNAs were transcribed using a MEGAshortscript T7 transcription kit (Thermo Fisher Scientific), purified using a MEGAclear Transcription Cleanup Kit (Thermo Fisher Scientific), and folded by thermal denaturation at 95°C for 5 min followed by equilibration at 37°C for 20 min in 10 mM Tris-HCl, pH 8.0. Reactivity of Exon 11 nucleotides to 1-methyl-7-nitroisatoic anhydride (1M7) was assessed by SHAPE-MaP using a variation of the standard protocol^24,25^, as follows.

RNA in negative control (−), modification reaction (+) and denatured control (dc) reactions were probed with the 1M7 acylation reagent under standard conditions. In separate reactions, the purified WT(−), WT(+), WT(dc), C1824U(−), C1824U(+), and C1824U(dc) RNAs were then reverse transcribed from a primer hybridized to nt 2139-2158 under conditions favoring mutagenesis instead of premature termination of reverse transcription, opposite acylated sites (50 mM Tris-HCl, pH 8; 75 mM KCl; 10 mM DTT; 500 uM dNTPs; 6 mM MnCl2; 200U SuperScript II Reverse Transcriptase; 5 pmol RNA; 2 pmol RT primer; 3 hr @ 42°C). Due to the lengths of the RNA segments being analyzed, the exon 11 sequences were divided into two overlapping zones (Z1: nt 913-1885, Z2: 1832-2047), and the respective zones in each of the six cDNA libraries amplified in separate 20-cycle PCR1 reactions (WT(−), Z1; WT(−), Z2, etc.; 12 reactions total) using appropriate primer sets containing partial Illumina adapter sequences. One-tenth of each PCR1 amplicon library was then reamplified in a 10-cycle PCR2 reaction using indexed primers suitable for library multiplexing and addition of the remaining Illumina adapter sequences. PCR2 amplicon libraries were purified from a 2% non-denaturing agarose gel, quantified using the KAPA Library Quantitation Kit, Illumina Platform, and sequenced on an Illumina MiniSeq using a Mid Output Kit (paired end, 2×150 nt).

Sequencing output was trimmed using a custom Python script to remove portions of each sequence read containing primer hybridization sites and to merge the respective forward and reverse Z1 and Z2 reads into individual FASTQ files (WT(−), WT(+), WT(dc), C1824U(−), C1824U(+), and C1824U(dc)). These files were processed using ShapeMapper software^24,25^ to produce 1M7 reactivity profiles that, together with the respective RNA sequences, were inputted into the RNA secondary structure prediction program RNAStructure^26^ to generate structure models. Images of the RNA structure models of lowest energy were refined using VARNA^27^.

### RNA isolation and RT-PCR analysis

RNA was isolated from cell culture samples at 72 hours post transfection using the RNeasy Plus Micro Kit (Qiagen). 1μg of each RNA sample was reverse-transcribed to cDNA using the iScript cDNA synthesis kit (Bio-Rad). RT-PCR was performed on the cDNA samples using the Quick-Load Taq 2X master mix (New England BioLabs). The primers used for RT-PCR are provided in **Supplementary Table 3**. DsRed expression was used as an internal loading control for the minigene experiments and TATA-binding protein (TBP) expression was used as a loading control for the ASO experiments. The RT-PCR protocol consisted of 30 seconds at 95°C, 30 cycles of 15 seconds at 95°C, 20 seconds at 55°C, and 30 seconds at 68°C, followed by a final extension of 5 minutes at 68°C. PCR products were resolved on a 2% agarose gel and imaged using a ChemiDoc MP Imaging System (Bio-Rad). Quantification of mRNA isoforms was performed using Image Lab 5.2.1 software (Bio-Rad).

### Quantitative PCR analysis

Quantitative PCR was performed on cDNA samples using the iQ SYBR Green Supermix (Bio-Rad) on a CFX96 Real Time PCR System (Bio-Rad) to measure LMNA and Δ150 expression levels. Primers used for qPCR are provided in **Supplementary Table 3**. The qPCR protocol consisted of 3 minutes at 95°C and 40 cycles of 20 seconds at 95°C and 30 seconds at 60°C. Reactions were performed in triplicate. Analysis was performed using the Bio-Rad CFX Maestro Software. LMNA and Δ150 expression levels were normalized to expression of TBP. Data are displayed as mean expression values ± SEM from a single experiment.

## Results

### SHAPE-MaP analysis of LMNA RNA secondary structure

LMNA pre-mRNA contains, in addition to its normal 5′ SS in exon 11, an internal alternative 5′ SS 150 nt upstream of the exon 11 3′ terminus (**Fig. 1A**), which is activated by the disease-causing C1824U point mutation in HGPS^20^. Usage of this alternative 5′ SS produces the Δ150 mRNA isoform, while usage of the normal 5′ SS produces the LMNA mRNA isoform (**Fig. 1A**). We exploited the presence of these competing 5′ SS as a model system for probing mechanisms of 5′ SS choice. To investigate the importance of RNA secondary structure in differential 5′ SS utilization, we used the chemical probing technology SHAPE-MaP to determine the secondary structures of WT and C1824U LMNA RNA constructs containing exons 10 and 11, and intron 11/12 (**Fig. 1A**).

**Figure 1.**
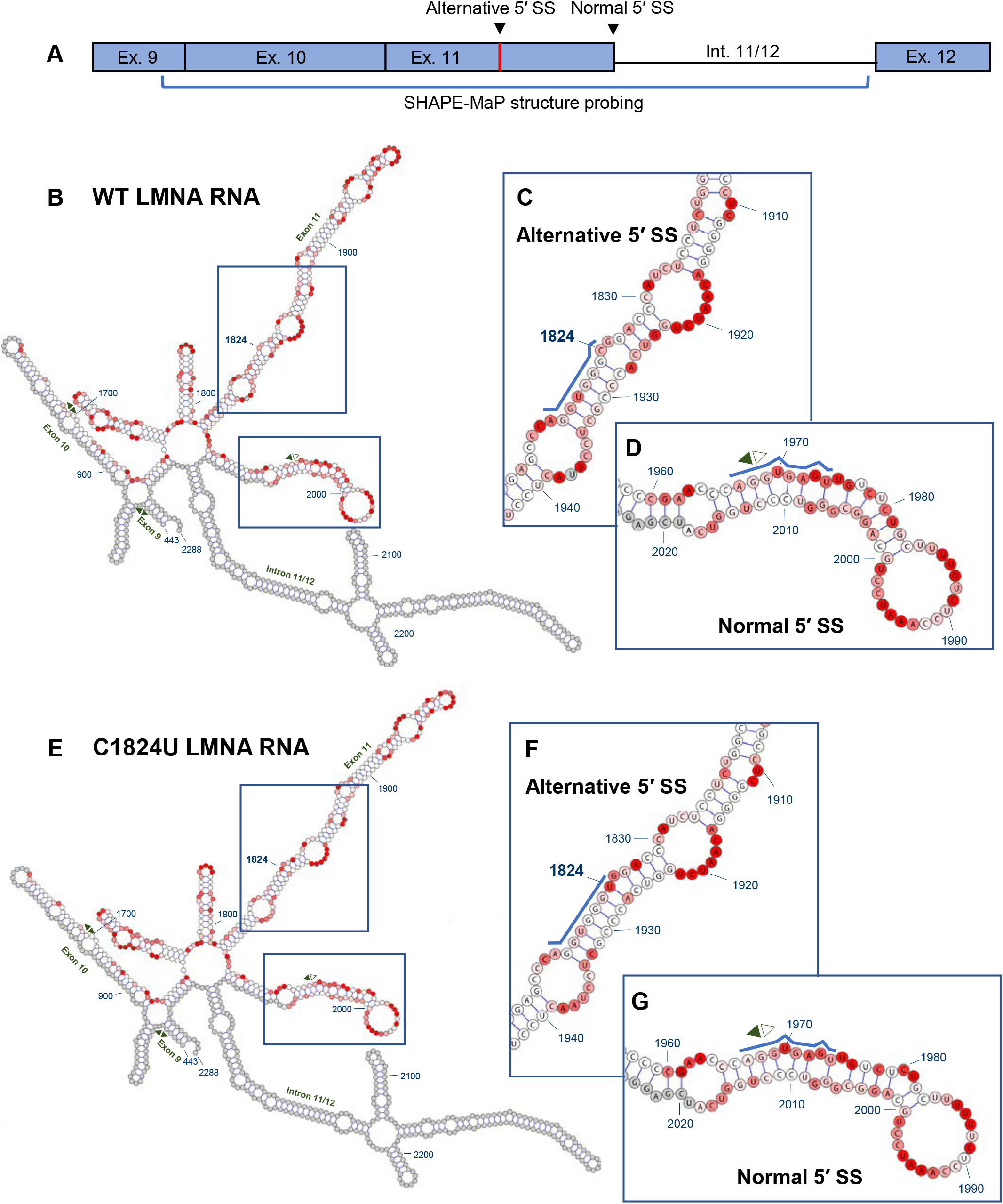
Secondary structures of wild-type (WT) and C1824U mutant LMNA RNA. RNA constructs probed by SHAPE-MaP were 687 nt in length and comprised the five 3′-terminal nt of exon 9, all of exons 10 and 11, and all but two nt of intron 11/12. Ribonucleotides are numbered in accordance with standard numbering of the human *LMNA* gene, and splice junctions are indicated by opposite-facing pairs of arrows (filled, toward exon; open, toward intron). Nucleotides for which no SHAPE-MaP data was obtained are shaded gray, while a red-to-white color gradient is used to indicate nucleotides that are more or less reactive to the 1M7 probing reagent, respectively. Within the insets, the alternative and normal 5′ SS sequences are highlighted with blue lines. **(A)** Schematic indicating the SHAPE-MaP structure probing region of LMNA. **(B)** WT LMNA construct secondary structure. **(C)** Sub-motif of WT LMNA RNA containing the alternative 5′ SS. **(D)** Sub-motif of WT LMNA RNA containing the normal 5′ SS. **(E)** C1824U mutant LMNA construct secondary structure. **(F)** Sub-motif of C1824U mutant LMNA RNA containing the alternative 5′ SS. **(G)** Sub-motif of C1824U mutant LMNA RNA containing the normal 5′ SS.

The WT LMNA RNA is highly structured, comprised of a series of interrupted stem-loop motifs arranged around a 6-way junction (**Fig. 1B**). The first and last of these stem-loop elements, principally comprising exon 10 and intron 11/12, respectively, are branched. A short 5′-to-3′ base pairing interaction is also predicted, suggesting that the entire construct may form a discrete, self-contained structure in the context of the larger LMNA RNA (**Fig. 1B**). The alternative exonic 5′ SS and the normal 5′ SS are contained in adjacent 142 nt and 80 nt hairpins, respectively, each interrupted by a series of bulges and internal loops. The former motif is stabilized by 9 consecutive base pairs between the junction and the first internal loop, while the corresponding segment of the latter is destabilized by a bulge and a mismatched pair. Importantly, the innermost 6 nt comprising the “core” of the alternative 5′ SS are predicted to be completely base paired, including four G-C pairs, three of which are consecutive (**Fig. 1C**). In contrast, although three guanines in the normal 5′ SS are also predicted to pair with cytosines, these G-C pairs are interrupted at position U1970, which represents one of two unpaired nucleotides in the normal 5′ SS core (**Fig. 1D**).

The structure of the C1824U mutant LMNA RNA closely resembles that of the WT LMNA RNA, indicating that the disease-causing mutation does not affect global LMNA RNA folding (**compare Fig. 1B and 1E**). Base pairing between U1824 and A1927 is predicted in the mutant RNA, although the impact of this on even local duplex stability is likely to be minimal (**Fig. 1F**). Imperfect pairing and high reactivity values within the distal regions of the normal 5′ SS hairpin produce slight differences in predicted local structures between the WT and HGPS mutant (**compare Fig. 1D with 1G**), suggesting that these nucleotides may be single stranded in alternative RNA conformations.

Taken together, these data show that the effect of the disease-causing C1824U mutation on global LMNA RNA structure is negligible but that the HGPS mutation results in a subtle local alteration of the alternative 5′ SS in exon 11. The structural mapping shows that the alternative 5′ SS nucleotides are sequestered in base pairing interactions within a long, stable RNA helix and that the normal 5′ SS exists in a less structured environment enabling it to sample alternative base pairing partners, including the U1-snRNA.

### Local structure around the point mutation is not sufficient to account for splice site activation

To determine how RNA secondary structure affects 5′ SS choice in exon 11 of LMNA, we conducted a systematic mutational analysis of predicted RNA structural features in a cell-based system (see Methods). Usage of the two 5′ SS was measured by RT-PCR with the long amplicon (LMNA) reflecting splicing at the normal 5′ SS and the short amplicon (Δ150) reflecting use of the exonic alternative 5′ SS (**Supplementary Fig. 1**; Methods).

Although SHAPE-MaP data suggests that the effects of the C1824U mutation on overall LMNA RNA structure are minimal, it is conceivable that the induced structural changes contribute to 5′ SS selection. To test this hypothesis, we introduced the more closed structure of the HGPS mutant into the WT background by generating mutants WT/1927A➔G and WT/1927A➔U, the latter of which re-created base pairing in the WT background of similar strength as in the mutant (**Fig. 2A, B**). However, neither of these mutants induced usage of the alternative 5′ SS (**Fig. 2D, E**). Conversely, introducing a more open structure, as present in the WT, into the region of the alternative 5′ SS in the mutant background (MUT/1928-1930C➔G; **Fig. 2C**), resulted in a slight increase in Δ150 expression and concomitant decrease in LMNA isoform expression (**Fig. 2D, E**). Overall, these results demonstrate that the local structural changes around the point mutation are not sufficient to account for the dramatic increase in usage of the alternative 5′ SS in HGPS.

**Figure 2.**
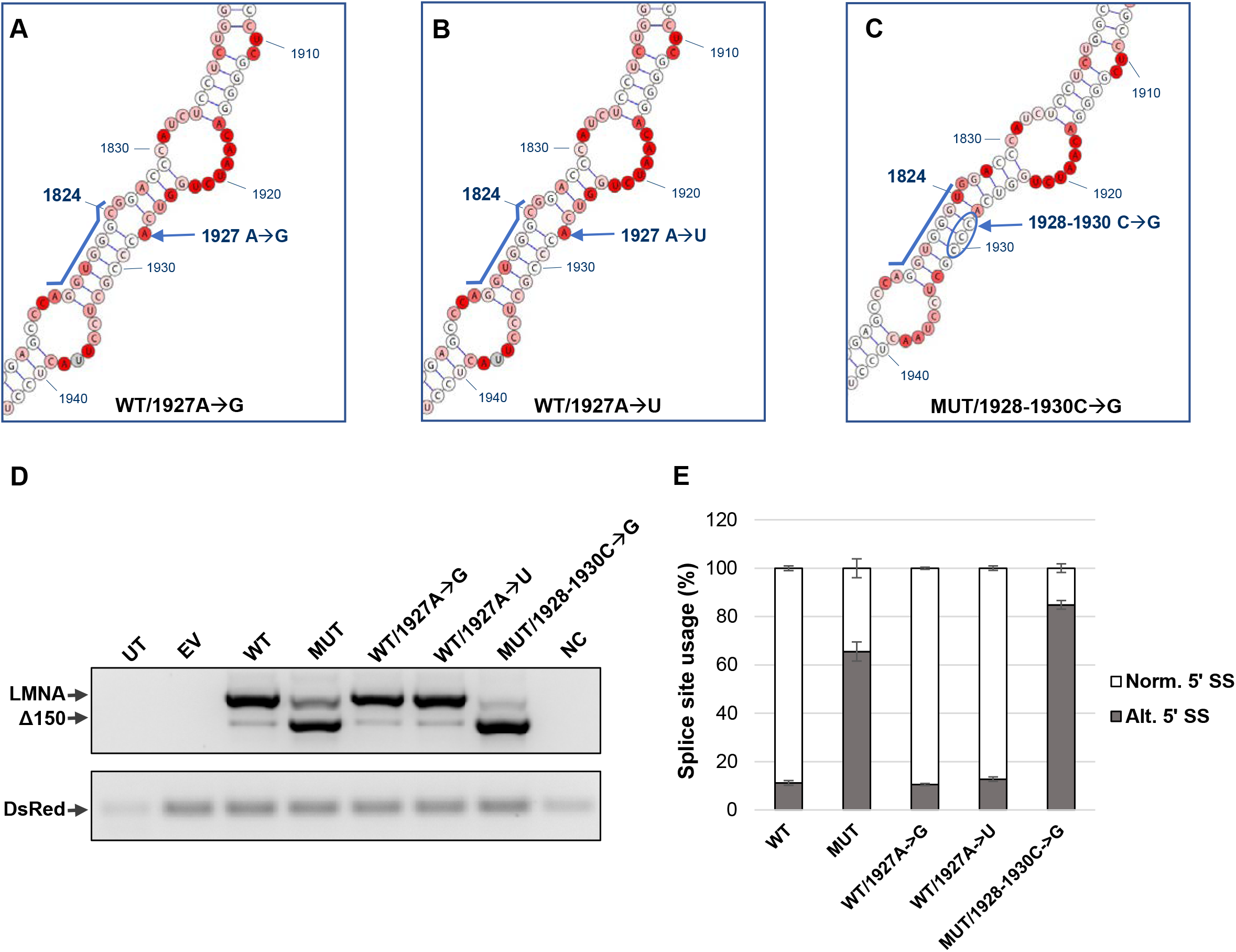
Local structural differences around the point mutation are not sufficient to account for alternative splice site usage in HGPS. **(A, B, C)** Design of minigene mutants. WT/1927A➔G **(A)** and WT/1927A➔U **(B)** were generated from the WT minigene construct. MUT/1928-1930C➔G **(C)** was generated from the MUT minigene construct. The alternative 5′ SS sequence is highlighted by a blue line. **(D)** RT-PCR analysis for LMNA/Δ150 expression levels of each mutant. 293T cells were transfected with 1μg of the indicated plasmids. RNA was harvested at 72 hours post transfection. DsRed expression served as an internal loading control. **(E)** Quantification of mRNA isoforms expressed as relative usage of the normal vs. alternative 5′ SS. Data are expressed as mean ± s.d. from three independent experiments. UT, untransfected; EV, empty vector; WT, LMNA wild-type minigene construct; MUT, LMNA C1824U mutant minigene construct; NC, negative control.

### Alternative 5′ SS usage is inhibited by base pairing with downstream elements

Based on the apparent increase in exonic alternative 5′ SS usage upon disruption of base pairing within the alternative 5′ SS of the MUT sequence, we sought to determine whether disrupting the alternative site hairpin in the WT background, where the alternative 5′ SS sequence is disfavored, would increase utilization of the site. Remarkably, introducing 1928-1930 triple C➔G mutations into the WT background (**Fig. 3A**) resulted in a pronounced increase in the amount of Δ150 product compared to WT (**Fig. 3D, E**), demonstrating that freeing the WT alternative 5′ SS from intrastrand base pairing substantially increases its usage. Disrupting the hairpin even further by mutating nucleotides 1928-1933 (**Fig. 3B**) resulted in an even greater increase in alternative 5′ SS usage relative to WT (**Fig. 3D, E**). This finding is in line with the almost complete suppression of aberrant LMNA RNA splicing observed in healthy individuals unaffected by HGPS^19^.

**Figure 3.**
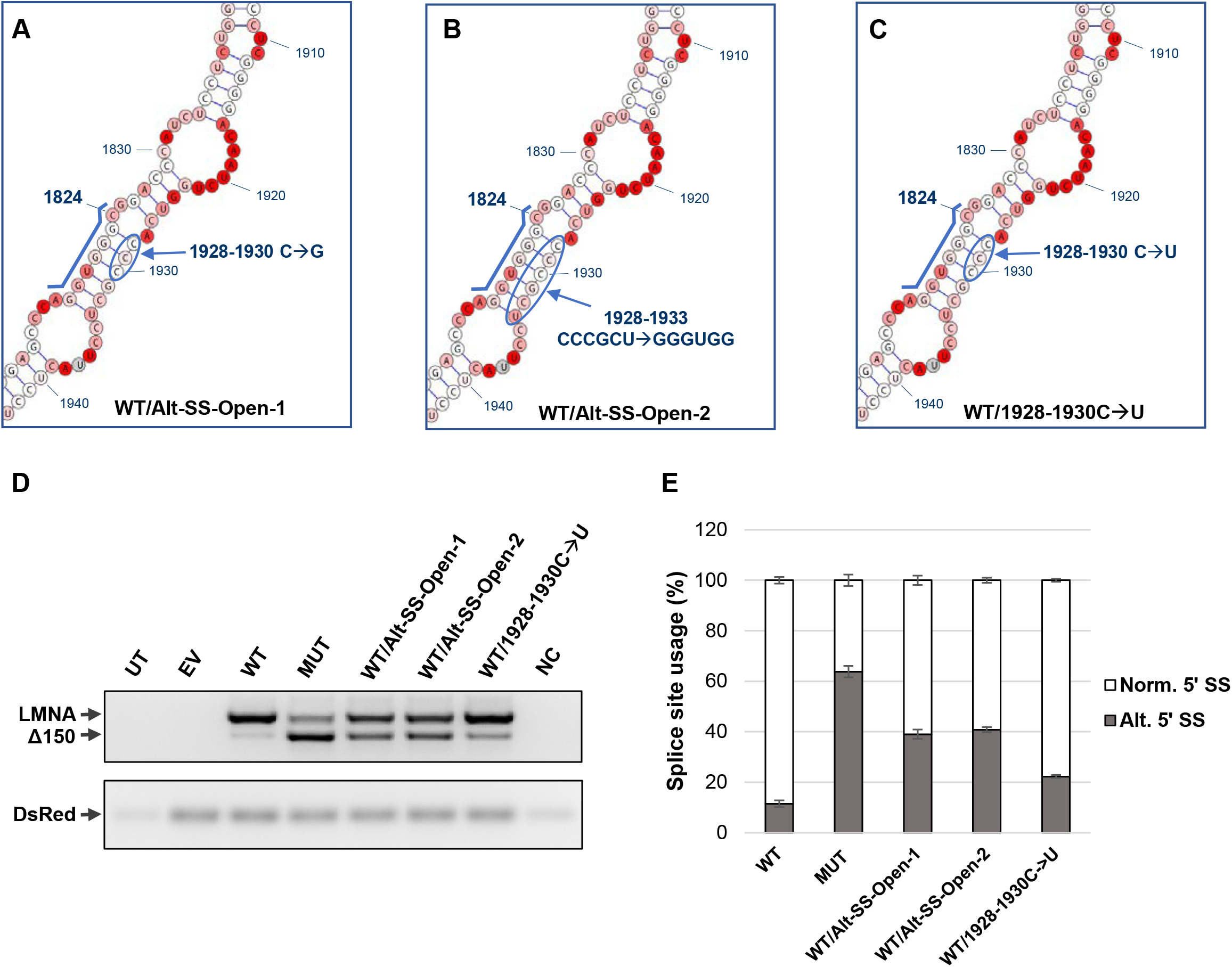
Splice site inhibition by downstream complementary sequence prevents usage of alternative splice site. **(A,B,C)** Design of the WT/Alt-SS-Open-1, WT/Alt-SS-Open-2, and WT/1928-1930C➔U mutants, generated from the WT minigene construct. The alternative 5′ SS sequence is highlighted by a blue line. **(D)** RT-PCR analysis for LMNA/Δ150 expression levels of each mutant. 293T cells were transfected with 1μg of the indicated plasmids. RNA was harvested at 72 hours post transfection. DsRed expression served as an internal loading control. **(E)** Quantification of mRNA isoforms expressed as relative usage of the normal vs. alternative 5′ SS. Data are expressed as mean ± s.d. from three independent experiments. UT, untransfected; EV, empty vector; WT, LMNA wild-type minigene construct; MUT, LMNA C1824U mutant minigene construct; NC, negative control.

Given these findings, we sought to confirm that the shift in splicing observed upon mutation of this downstream region was due to disruption of structural elements and not due to interference with sequence-based elements such as splicing factor binding sites. A triple C➔U mutation (**Fig. 3C**), which maintains the secondary structure at the alternative 5′ SS through wobble base pairing while changing the RNA sequence, resulted in a splicing pattern similar to the WT, with only a slight increase in alternative 5′ SS usage likely due to the weaker bonding strengths of the G-U vs. G-C base pairs (**Fig. 3D, E**).

To confirm that the effect of structural stability of the alternative 5′ SS observed in the minigene system reflects splicing outcome of endogenous LMNA, we designed an antisense oligonucleotide (ASO 1919) to hybridize to nucleotides 1919-1936 in the LMNA transcript and thus prevent intra-strand base pairing with the alternative 5′ SS (**Fig. 4A, B, green**). As a control, we used ASO 074 which hybridizes to nucleotides 1851-1868 in LMNA and has previously been shown to increase expression of the Δ150 isoform^28^ (**Fig. 4B, red**). In CRL 1474 normal human fibroblasts transfected with varying concentrations of ASOs, we observe a dose-dependent shift from predominant expression of the LMNA isoform to the Δ150 isoform at higher concentrations of ASO 1919, but not in the presence of the scrambled ASO (**Fig. 4C, D**). Taken together, these results suggest a mechanism by which alternative 5′ SS usage is suppressed by base paring with an RNA element ~100 nt downstream of the 5′ SS.

**Figure 4.**
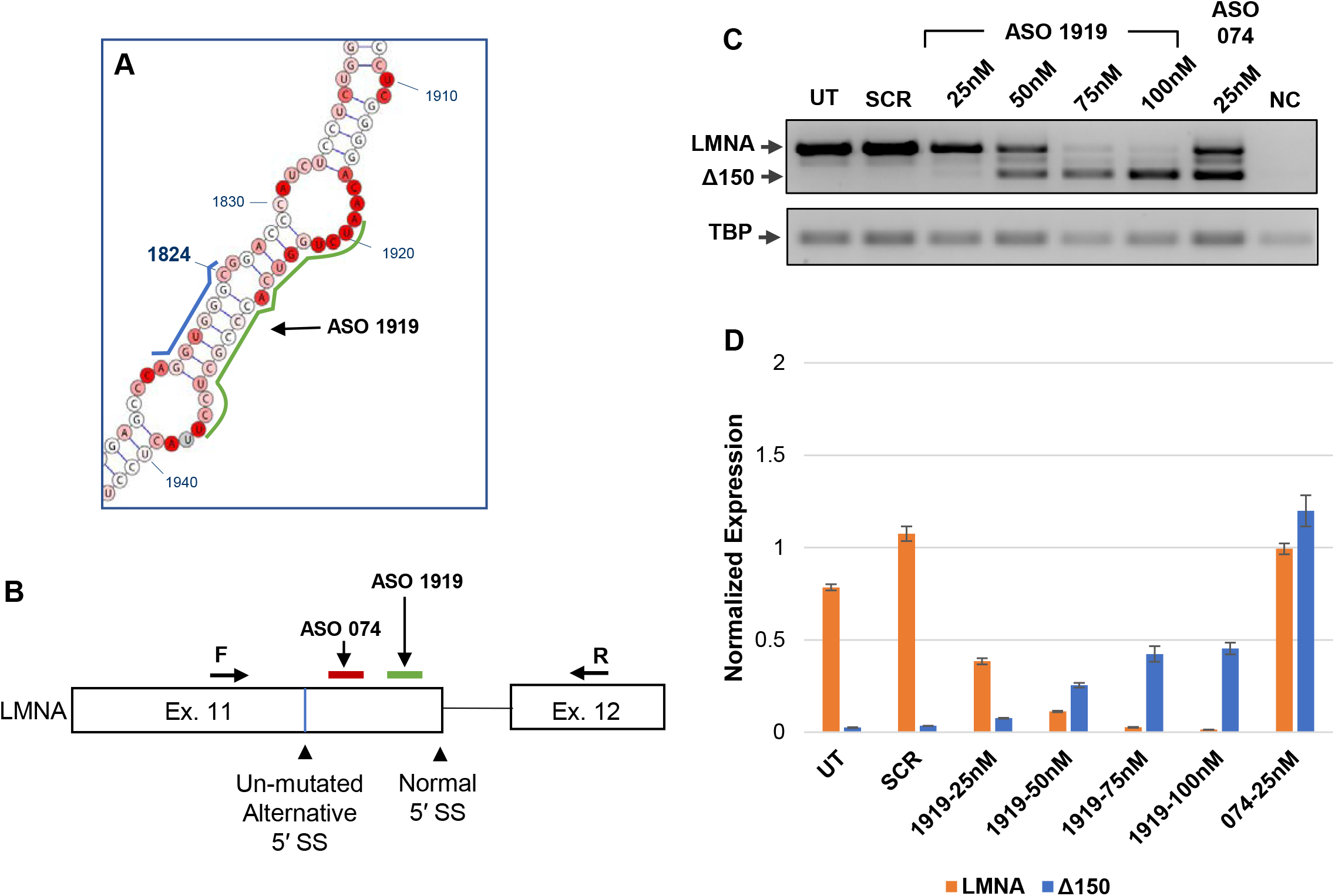
Increased usage of the endogenous alternative splice site in CRL 1474 cells following ASO treatment. **(A)** Diagram indicating the target location of ASO 1919 on the LMNA WT structural model, highlighted by a green line. The alternative 5′ SS sequence is highlighted by a blue line. **(B)** Diagram indicating the target location of ASO 074 and ASO 1919 in the context of the splicing cassette, highlighted by a red and green line, respectively. The locations of the forward and reverse primers used to measure endogenous expression of LMNA/Δ150 are indicated. **(C)** RT-PCR analysis for LMNA/Δ150 expression levels of each treatment group. CRL 1474 cells were transfected with 25-100nM of ASO 1919, 25nM of ASO 074, or 25nM of ASO SCR. RNA was harvested at 72 hours post transfection. TBP expression served as a loading control. **(D)** Quantitative PCR results for LMNA and Δ150 expression levels for the same samples tested in part **C**. Expression levels are normalized to expression of TBP. Data are expressed as mean ± SEM. ASO, antisense oligonucleotide; UT, untransfected; SCR, scrambled ASO; NC, negative control.

### Crosstalk between the 5′ SS affects splicing outcome

Given our observation of the importance of secondary structure in usage of the alternative exonic 5′ SS, we sought to determine whether usage of the normal 5′ SS is similarly affected by structural context. As a control, we first established that inactivation of the normal 5′ SS by mutation redirects splicing to usage of the exonic alternative 5′ SS (**Supplementary Fig. 2**). As expected, abolishing the normal 5′ SS in the WT construct by mutating it to a non-splice site sequence^10^ effectively repressed its usage (**Supplementary Fig. 2A, B, C**). We then generated two mutants from the WT construct in which the structural stability of the normal 5′ SS was increased by an intermediate or more substantive degree, respectively, through introduction of complementary base pairs (**Fig. 5A, B**). Both mutants led to increased usage of the alternative 5′ SS to an extent that exceeded that observed for WT LMNA, while usage of the normal 5′ SS was concomitantly reduced (**Fig. 5C, D**). Remarkably, stabilization of this region resulted in complete usage of the alternative 5′ SS (see WT/Norm-SS-Closed-2 mutant). Together, these findings support the notion that the structural stability of the normal 5′ SS, like that of the alternative 5′ SS, impacts splicing outcome.

**Figure 5.**
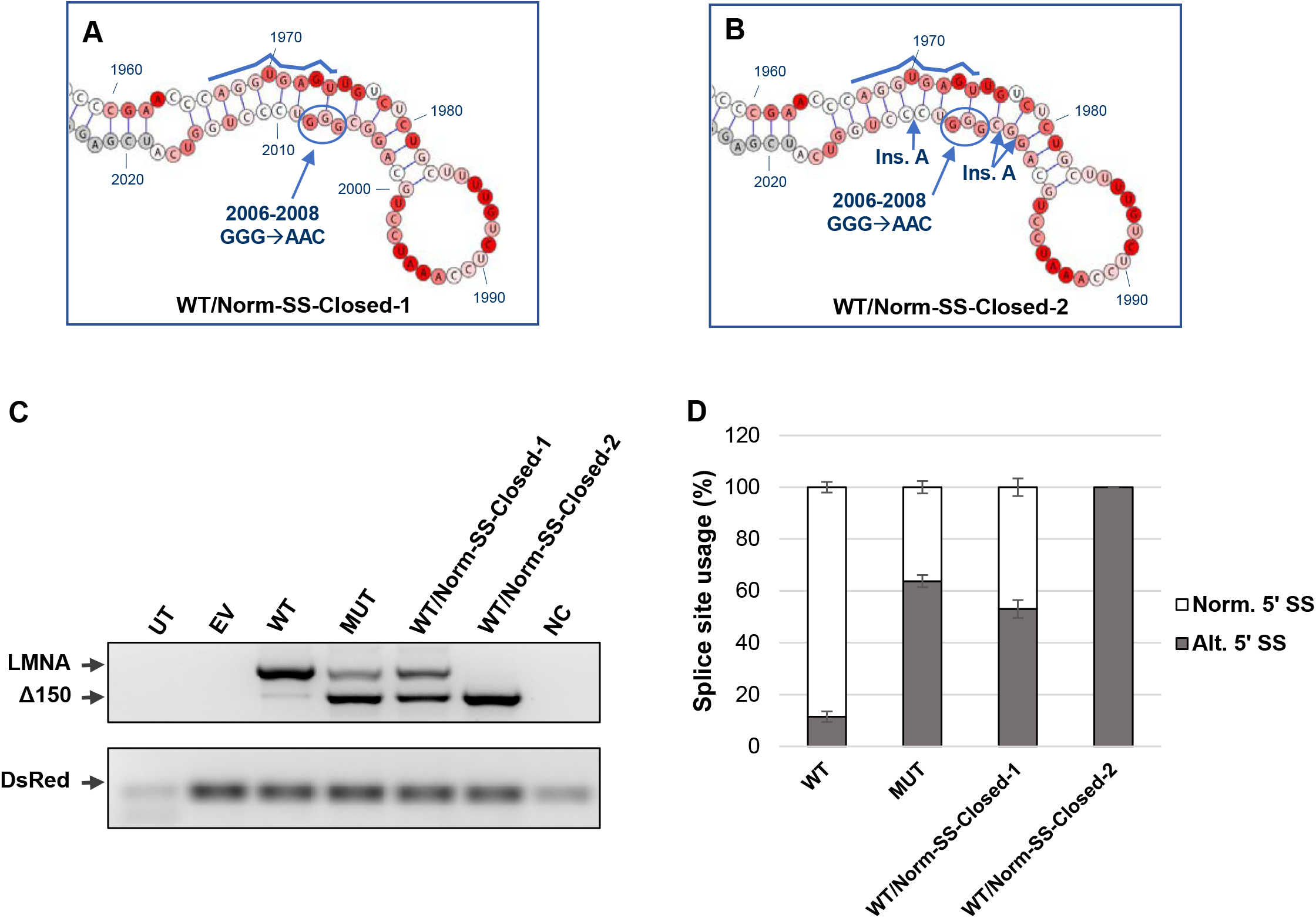
Increased structural stability around the normal splice site induces usage of the alternative splice site without the C1824U point mutation. **(A, B)** Design of the WT/Norm-SS-Closed-1 and WT/Norm-SS-Closed-2 mutants, generated from the WT minigene construct. The normal 5′ SS sequence is highlighted by a blue line. Sites at which adenines are inserted are indicated by ins. A. **(C)** RT-PCR analysis for LMNA/Δ150 expression levels of each mutant. 293T cells were transfected with 1μg of the indicated plasmids. RNA was harvested at 72 hours post transfection. DsRed expression served as an internal loading control. **(D)** Quantification of mRNA isoforms expressed as relative usage of the normal vs. alternative 5′ SS. Data are expressed as mean ± s.d. from three independent experiments. UT, untransfected; EV, empty vector; WT, LMNA wild-type minigene construct; MUT, LMNA C1824U mutant minigene construct; NC, negative control.

### The HGPS point mutation confers a dominant effect on 5′ SS choice

In part due to its potential relevance to designing HGPS therapeutics, we next asked whether altering the structural stability of the alternative and normal 5′ SS may redirect splicing in the MUT minigene construct towards expression of the WT LMNA isoform. To do so, we designed three C1824U mutant variants to either further stabilize the stem containing the alternative site, or alternatively to destabilize the stem containing the normal site or to combine these two mutations (**Supplementary Fig. 3A, B**). In all three mutants, regardless of the structural stability of the available 5′ splice sites, the alternative site is always favored (**Supplementary Fig. 3C, D**). We thus conclude that the HGPS point mutation confers a dominant effect on 5′ SS choice in LMNA.

### Sequence-independent preferential use of the distal 5′ SS

Our finding that the mutant alternative 5′ SS in LMNA is strongly favored regardless of changes in structural stability makes it clear that other factors contribute to its preferred usage. In addition to primary sequence and structural organization, we expected a further contributor to 5′ SS choice to be the relative position of competing splice sites in the primary RNA sequence. To assess the contribution of location to splice site choice, we generated a series of mutants containing two normal 5′ SS, two mutated (1824U) alternative 5′ SS, or two unmutated (1824C) alternative 5′ SS sequences in the proximal and distal 5′ SS positions (**Fig. 6A**). Additionally, we created a mutant in which the sequence of the normal 5′ SS was swapped with that of the 1824U alternative 5′ SS (**Fig. 6A**). We found that when the proximal and distal splice site positions in exon 11 of LMNA RNA have identical sequence compositions, the distal position is always predominantly used as indicated by greater relative abundance of the Δ150 product in all mutants (**Fig. 6B, C**). The preference for alternative 5′ SS usage was even more pronounced for the SWAP mutant, likely because both sequence and position favor its use (**Fig. 6B, C**). We conclude that relative positioning of competing 5′ SS is a critical determinant of splice site choice in HGPS.

**Figure 6.**
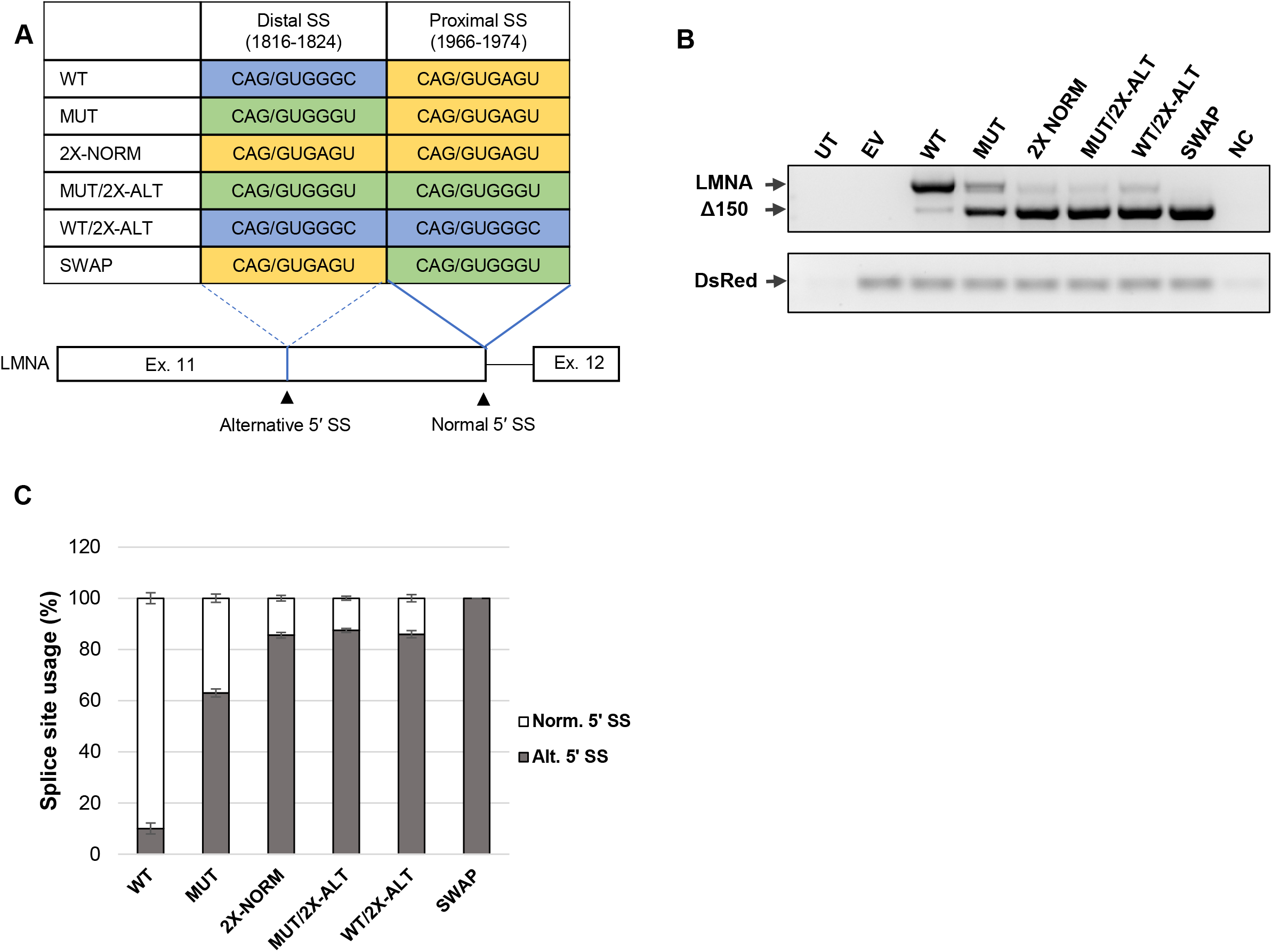
Positional preference for the distal splice site over the proximal splice site during splice site selection. **(A)** Design of mutant minigene constructs, comparing sequences used for the proximal and distal splice site locations. Locations indicated are based on the numbering of the genomic sequence. Proximal and distal terms used in reference to intron 11/12. Forward slash indicates the splice junction. Splice sites with the same color have the same sequence. **(B)** RT-PCR analysis for LMNA/Δ150 expression levels of each mutant. 293T cells were transfected with 1μg of the indicated plasmids. RNA was harvested at 72 hours post transfection. DsRed expression served as an internal loading control. **(C)** Quantification of mRNA isoforms expressed as relative usage of the normal vs. alternative 5′ SS. Data are expressed as mean ± s.d. from three independent experiments. WT, LMNA wild-type minigene construct; MUT, LMNA C1824U mutant minigene construct; 2X-NORM, minigene construct with normal splice site sequence at both proximal and distal splice site positions; MUT/2X-ALT, minigene construct with C1824U mutant alternative splice site sequence at both proximal and distal splice site positions; WT/2X-ALT, minigene construct with un-mutated alternative splice sequence at both proximal and distal splice positions; SWAP, minigene construct with C1824U mutant alternative splice site sequence at the proximal splice site and normal splice site sequence at the distal splice site; NC, negative control.

## Discussion

Sequence analysis alone frequently fails to accurately predict experimentally determined splicing outcomes and splice site usage^14^, strongly suggesting that other factors contribute to splice site choice. Here we have used a combination of SHAPE-MaP RNA secondary structural analysis and targeted mutagenesis to investigate the contributions of RNA sequence, RNA secondary structure, distal regulatory elements and linear position to 5′ SS choice. Using the disease-relevant alternative splicing of LMNA in HGPS as a model system, we have revealed an intricate interplay among RNA structure, linear position, and sequence in 5′ SS selection.

Our SHAPE-MaP analysis describes the complex higher order structure of the LMNA pre-mRNA. Interestingly, only minor local differences between the wild-type and mutant forms of LMNA pre-mRNA were found, despite the fact that they undergo distinct alternative splicing fates, with dramatic cellular and organismal consequences, due to differential 5′ SS usage. The structural models for wild-type and mutant LMNA obtained here differ somewhat from previously published structures^22^, most likely reflecting the fact that the RNA constructs previously probed are markedly shorter than those utilized here (329 nt vs. 687 nt), precluding detection of long-range interactions that are critical for the overall structure of the RNA. Our combined use of SHAPE-MaP and RNAStructure software is also advantageous in that the 1M7 probing reagent can acylate all four nucleotides, problems with steric clashes that can sometimes alter enzymatic cleavage profiles are eliminated, and 1M7 reactivity values can be incorporated as soft, pseudoenergy constraints into the RNAStructure folding algorithm, from which unbiased models that account for all of the probing data can be generated^29^. By harnessing the advantages of SHAPE-MaP, we were able to characterize the structure of LMNA RNA and elucidate the relationship of RNA structure to alternative splicing fate.

We took advantage of the presence of two competing 5′ SS in the LMNA pre-mRNA to probe the contribution of structural features to 5′ SS selection. We determined that structural features of LMNA RNA identified by SHAPE-MaP were important determinants of 5′ SS selection in vivo. Specifically, we established that a complementary region ~100 nt downstream of the alternative exonic 5′ SS serves to sterically hinder its accessibility to the splicing machinery. Disruption of base pairing between this region and the alternative 5′ SS facilitates its selection, demonstrating the inhibitory function of the downstream element. This finding is supported by preferential usage of the alternative 5′ SS lacking the HGPS point mutation upon inactivation of the normal 5′ SS. Importantly, we also find that increasing the rigidity of the local structure at a splice site diminishes its selection and promotes usage of competing sites. These findings extend previous studies that show evidence for the influence of secondary structures in 5′ SS suppression in other systems, suggesting that this may be an important general mechanism of alternative splicing regulation^15,30,31^.

In addition to demonstrating a role of RNA structural stability on 5′ SS choice, we also identify relative positioning in the primary RNA sequence as a key determinant in 5′ SS selection. We observe a prominent advantage of the distal 5′ SS over the proximal 5′ SS when the two positions are composed of identical sequences, regardless of the sequence of the splice site. It appears that this advantage may not be universal, however, as other studies report an advantage for proximal 5′ SS selection^10–13^. A possible explanation is the use of different model systems and the context of the candidate sites, highlighting the need to consider multiple parameters when assessing mechanisms of 5′ SS selection.

Given the disease relevance of the model system used here, our findings have implications for HGPS. Based on our observations, it seems likely that a therapeutic strategy aimed at switching 5′ SS selection from the alternative 5′ SS to the normal 5′ SS by stabilizing the alternative splice site and making the normal splice site more open will not be successful due to the strong effect of the alternative 5′ SS location along the transcript, as observed here. Our data indicate that in WT LMNA RNA, the positional advantage of the alternative 5′ SS is not sufficient to outcompete the structural and sequence-based advantages favoring the normal 5′ SS (**Fig. 7**). However, the disease-causing mutation sufficiently enhances the strength of the alternative 5′ SS, so that, combined with its positional advantage, it outweighs the sequence-based advantage of the normal 5′ SS (**Fig. 7**). Experimentally, this effect is mimicked by disrupting the structure by either reducing the stability of the alternative 5′ SS or increasing the stability of the normal 5′ SS, tipping the balance towards usage of the alternative 5′ SS (**Fig. 7**).

**Figure 7.**
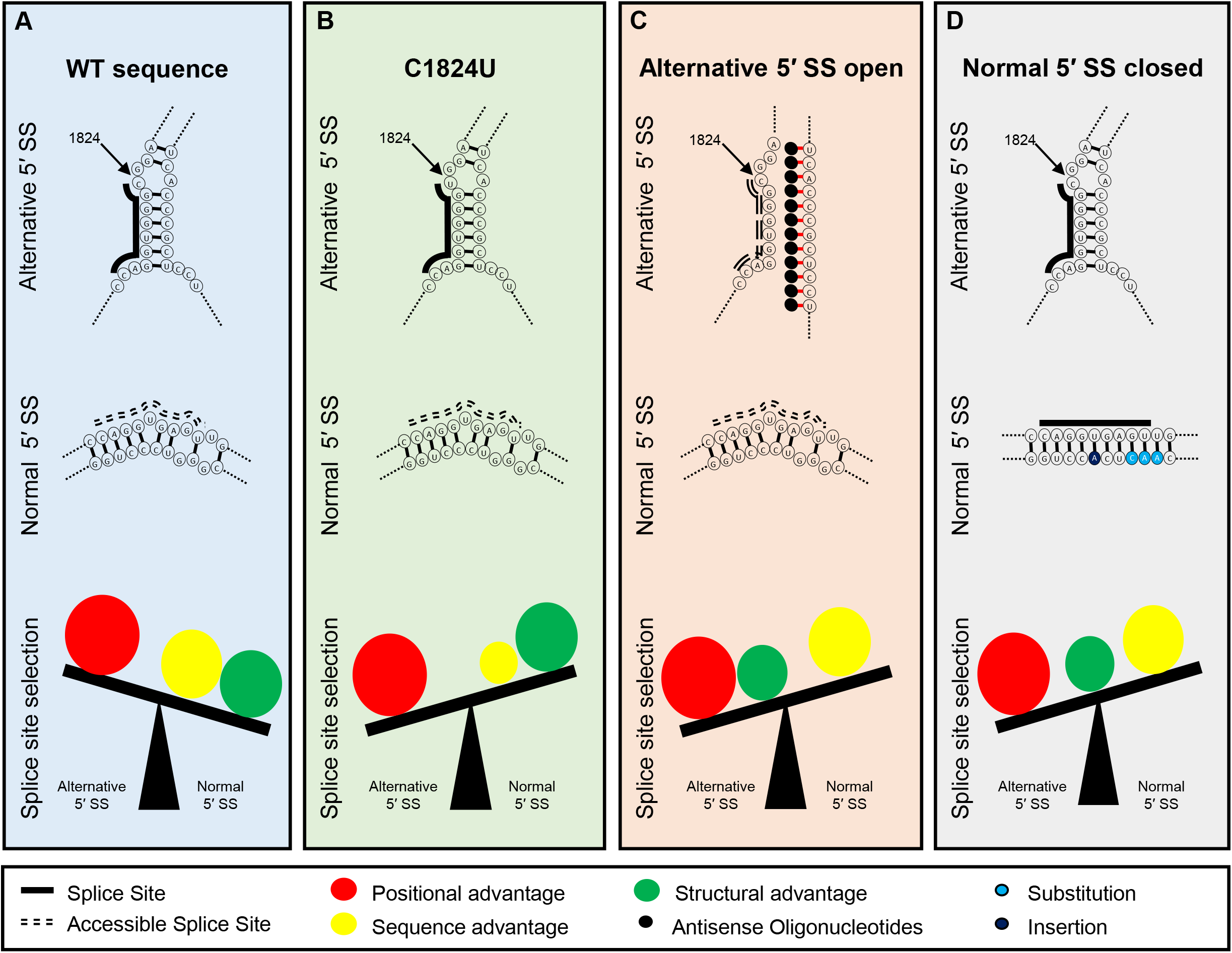
A model for combinatorial elements that orchestrate 5′ SS selection in exon 11 of LMNA. **(A)** In the wild-type sequence, usage of the alternative 5′ SS in hindered by stable base pairing with the downstream sequence. Thus, it is not as accessible as the normal 5′ SS which forms an unstable structure. While the positional advantage favors the alternative 5′ SS, the sequence and structure favor selection of the normal 5′ SS. **(B)** In the case of C1824U mutation, the sequence advantage for the normal 5′ SS is not as potent as in the wild-type sequence. Thus, the positional advantage of the alternative 5′ SS shifts the balance to favor its selection. **(C)** ASO treatment interfering with the inhibitory structural elements of the alternative splice site confers the structural advantage to the alternative 5′ SS. Combined with the positional advantage, the alternative 5′ SS is selected even though the sequence favors the normal 5′ SS. **(D)** Reducing reactivity of the normal 5′ SS, by making the region more structurally stable, eliminates the structural advantage of the normal 5′ SS. The alternative 5′ SS is selected due to the positional advantage over the sequence advantage of the normal 5′ SS.

In summary, the data provided here offers a novel perspective on alternative 5′ splice site selection. While this study has clarified the influence of sequence, structure and position in 5′ SS selection of exon 11 of LMNA, the generalizability of these findings need to be confirmed experimentally for other spliced RNAs. Additionally, there are unanswered questions regarding the transition from 5′ SS recognition to selection and whether a threshold of 5′ SS/U1 binding strength is required for splice site commitment. Furthermore, the influence of other splicing factors and the implications of RNA polymerase II transcription kinetics on splice site choice require further characterization. Regardless, our observations highlight the involvement of multiple factors in 5′ SS selection and suggest that consideration of parameters other than sequence, particularly secondary RNA structure, is warranted when predicting 5′ SS usage.

## Acknowledgements

This work was supported by the Intramural Research Program of the National Institutes of Health (NIH), National Cancer Institute, and Center for Cancer Research.

## Author Contributions

FT, AS, TM and SG conceived the study, FT, AS, and JR performed all experiments, and TM and SG supervised the study. All authors contributed to data analysis and interpretation, and writing of the manuscript.

